# Immature leaves are the dominant volatile sensing organs of maize

**DOI:** 10.1101/2023.01.12.523648

**Authors:** Lei Wang, Simon Jäggi, Mario Walthert, Jamie M. Waterman, Tristan M. Cofer, Matthias Erb

## Abstract

Plants perceive herbivory induced volatiles and respond to them by upregulating their defenses. So far, the organs responsible for volatile perception remain poorly described. Here, we show that responsiveness to the herbivory induced green leaf volatile (*Z*)-3-hexenyl acetate (HAC) in terms of volatile emission, transcriptional regulation and defense hormone activation is largely constrained to younger maize leaves. Older leaves are much less sensitive to HAC. In a given leaf, responsiveness to HAC is high at immature developmental stages and drops off rapidly during maturation. Responsiveness to the non-volatile elicitor ZmPep3 shows an opposite pattern, demonstrating that hyposmia is not driven by defective canonical defense signaling. Neither stomatal conductance nor leaf cuticle composition explain the unresponsiveness of older leaves to HAC, suggesting perception mechanisms upstream of canonical defense signaling as driving factors. Finally, we show that hyposmia in older leaves is not restricted to HAC, and extends to the full blend of herbivory induced volatiles. In conclusion, our work identifies immature maize leaves as dominant stress volatile sensing organs. The tight spatiotemporal control of volatile perception may facilitate within-plant defense signaling to protect young leaves, and may allow plants with complex architectures to explore the dynamic odor landscapes at the outer periphery of their shoots.

## Introduction

Plants can sense various herbivory-related cues to initiate defense. These danger cues can be derived from herbivores or damaged plant tissues (Erb and Reymond, 2019; Reymond, 2021; Tanaka and Heil, 2021). Danger cues can be translocated as signals via the vasculature or as volatiles via the air The perception of danger signals then triggers anti-herbivore defenses via activation of the jasmonate signaling pathway (Howe and Jander, 2008; Erb et al., 2012). Recent years have seen considerable advances in understanding herbivory-derived cues as well as vascular and volatile defense signaling (Hu et al., 2019; Ye et al., 2019; Steinbrenner et al., 2020; Poretsky et al., 2021). However, how plants perceive volatile danger cues is still poorly understood.

Green leaf volatiles (GLVs) are C6 aldehydes, alcohols, or esters. These membrane lipid-derived volatiles are rapidly produced and emitted upon leaf damage caused from mechanical wounding or insect feeding GLV responsiveness is common across many plants (Ameye et al., 2018; Meents and Mithöfer, 2020; Wang and Erb, 2022). GLVs trigger defense responses in maize, *Arabidopsis*, tomato, and poplar (Engelberth et al., 2004; Frost et al., 2008; Asai et al., 2009). In maize, exposure to the GLV (*Z*)-3-hexenyl acetate (HAC) induces JA biosynthesis, the production of benzoxazinoid secondary metabolites, and the emission of volatiles in undamaged plants (Engelberth et al., 2004; Hu et al., 2019). HAC-induced volatiles in maize include indole, monoterpenes, sesquiterpenes and the homoterpenes (*E*)-4,8–dimethyl–1,3,7-nonatriene (DMNT) and *(E,E*)-4,8,12-trimethyltrideca-1,3,7,11-tetraene (TMTT) (Engelberth et al., 2004). Other GLVs, such as (*Z*)-3-hexenal and (*Z*)-3-hexen-1-ol, trigger similar defense responses in maize (Engelberth et al., 2004).

Volatile sensing likely requires direct access of volatiles to the plasma membrane or intracellular space. Little is known about how volatiles enter plant leaves. However, several routes of volatile exchange between plant tissues and the atmosphere have been described (Liao et al., 2021; Lin et al., 2021; Wang and Erb, 2022; Widhalm et al., 2022). In maize, tomato and soybeans, stomata control volatile release (Seidl-Adams et al., 2015; Lin et al., 2021). In petunia flowers, the hydrophobic cuticle acts both as a resistance barrier and a sink of volatiles. Its thickness thus regulates the emission of volatiles (Liao et al., 2021). Leaf cuticular waxes can also be a volatile sink for exogenous volatiles (Camacho-Coronel et al., 2020). Volatile perception itself may be mediated by interactions of volatiles with the plasma membranes or by intra- or extracellular volatile-binding proteins (Loreto and D’Auria, 2022; Wang and Erb, 2022). Thus, the current scenario of volatile uptake and perception is that volatiles diffuse into the leaf apoplast via the stomata and the cuticle. They may then interact with the plasma membrane or membrane-bound proteins, or be transported into leaf cells to interact with intracellular receptors (Wang and Erb, 2022). So far, the importance of stomata and the leaf-cuticle as gateways for volatile perception is unclear (Widhalm et al., 2022).

The perception of environmental and danger cues by plants is strongly regulated by developmental processes. In roots, gravity sensing, perception of nitrate and phosphorus are confined to the root cap at the root tip (Svistoonoff et al., 2007; O’Brien et al., 2016; Su et al., 2017). In *Arabidopsis thaliana*, perception of the microbe-associated molecular pattern flg22 in the rhizosphere also happens mainly at the root cap, extending to the transition/elongation zone, but is absent in differentiated root parts (Zhou et al., 2020). In *Nicotiana benthamiana*, responses to bacterial cold shock proteins (CSPs) are high in the leaves of flowering plants, but weaker or even absent in very young plants (Saur et al., 2016; Wang et al., 2016). Conversely, in *Nicotiana attenuata,* the herbivory-induced accumulation of defense hormones decreases as plants reach the flowering stage (Diezel et al., 2011). As both volatile uptake and sensing are likely also regulated by developmental processes, volatile perception by plants may involve distinct tissues and/or growth stages (Wang and Erb, 2022). Aboveground parts that specifically take up and perceive volatiles could act as volatile sensing organs (Wang and Erb, 2022).

Here, we investigate the capacity of different maize leaves at different developmental stages to respond to herbivory induced volatiles such as HAC. We compare responses to HAC and non-volatile danger cues to determine whether volatile sensitivity is a specific feature of developing leaves. We track HAC responsiveness in real time by measuring the induction of aromatic and terpenoid volatiles following contact with HAC and non-volatile defense cues. We use transcriptional and hormonal profiling to determine at which level of defense signaling the differences in responsiveness originate. Finally, we perform experiments with cuticular wax mutants and measure stomatal conductance to determine whether the differences in HAC responsiveness can be explained by changes in uptake due to differences in cuticular or stomatal development. Finally, we perform experiments with full blends of herbivory induced plant volatiles to determine whether differences in responsiveness also govern responses to natural combinations of bioactive volatiles. Taken together, our results show that immature leaves are the dominant volatile sensing organs of maize plants.

## Results

### Mature maize leaves are largely unresponsive to (*Z*)-3-hexenyl acetate

Maize plants have been shown to release volatiles in response to herbivory-induced volatiles. Green leaf volatiles such as (*Z*)-3-hexenyl acetate (HAC) were identified to mediate this process. We confirmed this finding using intact, detached V4 maize seedlings that were exposed to dispensers releasing HAC at a physiological dose of ∼70 ng*h^-1^. Using a newly developed high-throughput, high resolution PTR-MS (proton transfer reaction-mass spectrometry) system (Krechmer et al., 2018), we found that seedlings started to release significant amounts of indole, monoterpenes, sesquiterpenes and the homoterpenes DMNT and TMTT one hour after HAC exposure (Fig. 1A). Induced emissions dropped during the transition to the night phase. As soon as lights were turned on the next day, HAC-induced volatile emissions resumed, albeit at lower levels. To test whether single leaves are able to respond to HAC, we excised mature L4 leaves of V4 seedlings and measured HAC-induced volatile release. The leaf base was always submerged in water to avoid drought stress and stomatal closure. Surprisingly, we found no induction of terpenes, and only a minor induction of indole (Fig. 1B). A similar pattern was observed when L4 leaves were excised together with the leaf sheath (Supp. Fig. 2). To test whether this lack of responsiveness is due to a defect in canonical defense signaling and/or an incapacity of L4 leaves to release volatiles, we treated the leaves with the (non-volatile) peptide ZmPep3, a known inducer of jasmonate defense signaling and volatile release in maize (Huffaker et al., 2013). ZmPep3 treatment induced a strong and persistent release of all measured volatiles in detached L4 leaves (Supp. Fig. 1). These results show that mature L4 leaves are largely unresponsive to HAC, despite their intact and highly responsive defense signaling and volatile biosynthesis pathways.

**Figure 1.**
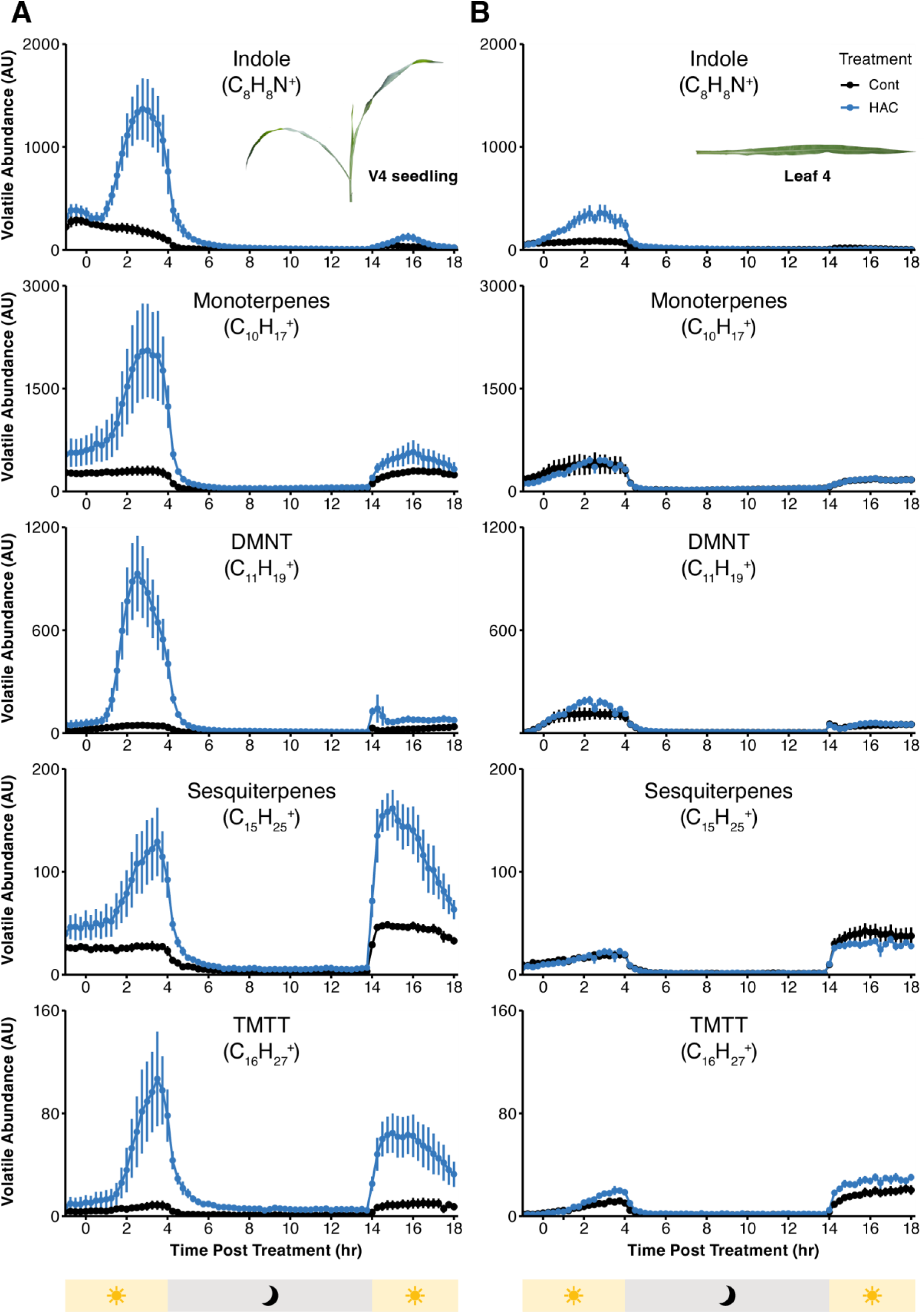
Mature maize leaves are largely unresponsive to HAC. A) Volatile emission of detached V4 seedlings exposed to HAC dispensers (release rate: 70 ng*h^-1^). B) Volatile emission of detached L4 leaves of V4 seedlings exposed to HAC dispensers. Untreated seedlings and leaves were used as controls (Cont). Distinguished by their exact mass, relative emission rates of indole, monoterpenes, DMNT, sesquiterpenes and TMTT were measured by PTR-MS. Light and dark periods are indicated below the X-axis. Data are shown as mean ± s. e. (n = 4 for V4 seedlings, n = 6 for L4 leaves).

**Supplementary Figure 1.**
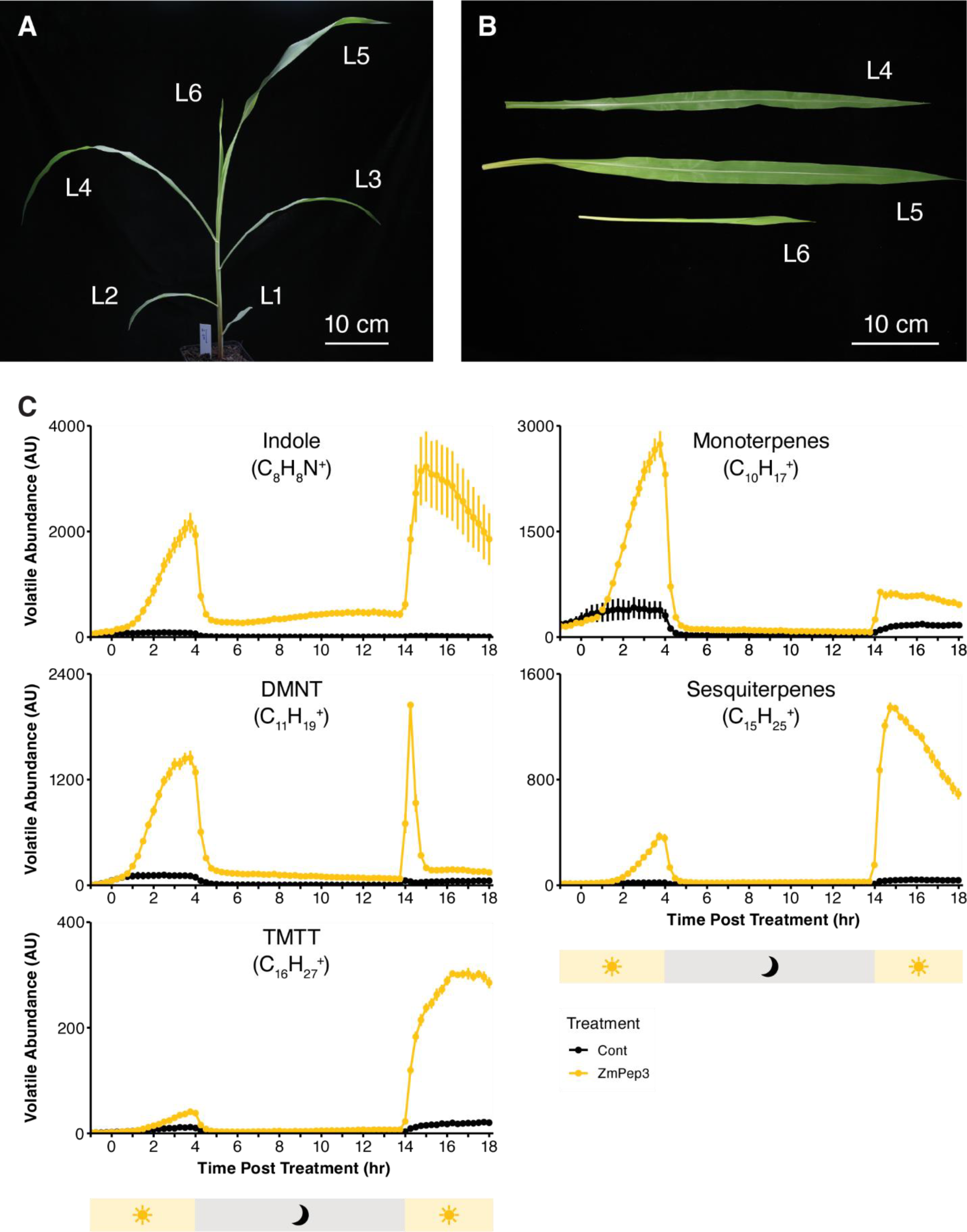
ZmPep3-induced VOCs emission in detached L4 leaves. A) Picture of a V4 stage B73 maize seedling. Leaves are numbered in ontogenetical order. B) Picture of detached L4, L5 and L6 leaves from a V4 maize seedling. C) ZmPep3-triggered volatile emission in detached L4 leaves from V4 seedlings. Detached L4 leaves were treated with 1 µM ZmPep3. Untreated leaves were included as controls. Distinguished by their exact mass, five groups of volatiles were analyzed: indole, monoterpenes, DMNT, sesquiterpenes and TMTT. Yellow bar below the x-axis indicates VOCs emission under light, grey bar indicates VOCs emission in the dark (4 – 14 h post treatment). Data are shown as mean ± s. e. (n = 6).

**Supplementary Figure 2.**
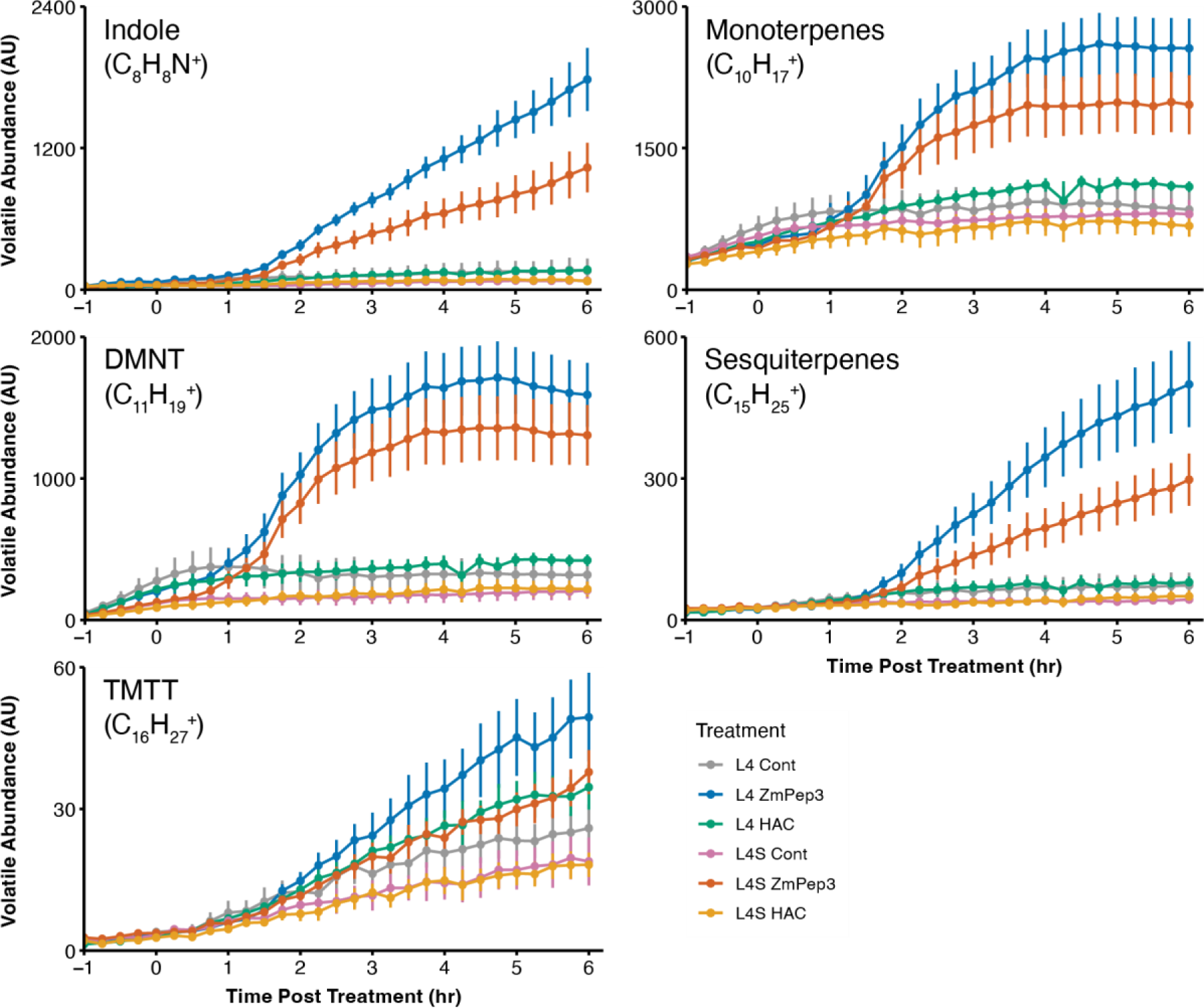
HAC-induced VOCs emission in L4 leaves of V4 seedlings. Detached L4 leaves (with or without leaf sheath) from V4 maize seedlings were treated with 1 µM ZmPep3 or HAC dispenser. L4: Leaf 4 without sheath (leaf blade only); L4S: Leaf 4 with 6 cm sheath. Distinguished by their exact mass, relative emission rates of indole, monoterpenes, DMNT, sesquiterpenes and TMTT were measured by PTR-MS for 6 hours after treatment. Data are shown as mean ± s. e. (n = 6).

### Volatile responsiveness to (Z)-3-hexenyl acetate is strongest in young maize leaves

Based on our results with detached L4 leaves, we hypothesized that within a maize seedling, HAC responsiveness may be conferred by younger leaves. We thus dissected L4, L5 and L6 leaves from V4 seedlings and exposed them to HAC individually (Fig. 1B). We normalized the volatile emission to leaf fresh weight to account for the substantial differences in leaf biomass of the different leaves. We also included ZmPep3 treatments as controls. We observed a marked shift from strong ZmPep3 responsiveness in L4 leaves to strong HAC responsiveness in L6 leaves. L5 leaves showed an intermediate phenotype, responding equally to both danger cues (Fig. 2, Supp. Table. 1). To further confirm that the differential volatile emission in L4, L5 and L6 leaves is specific to elicitation by HAC, we also used *Spodoptera exigua* (*S. exigua*) simulated herbivory to induce individual leaves. The treatment consisted of wounding the leaves and applying *S. exigua* oral secretions, thus including the full blend of non-volatile herbivory-associated molecular patterns (Erb et al., 2015). Similar to elicitation with ZmPep3, L4 leaves showed the highest emission of indole, monoterpenes, DMNT and sesquiterpenes, and responsiveness dropped in L5 and L6 leaves (Supp. Fig. 3). A notable exception was TMTT emission, which showed the opposite pattern in response to oral secretions than to ZmPep3 treatment (Supp. Fig. 3). In a separate experiment, we tested if leaf three (L3), an even older leaf, is responsive to HAC exposure. Similar to L4, its volatile emission after HAC exposure was comparable to the untreated leaf (Supp. Fig. 4). Together, these experiments show that HAC volatile responsiveness is specifically restricted to young maize leaves, with older leaves becoming unresponsive.

**Figure 2.**
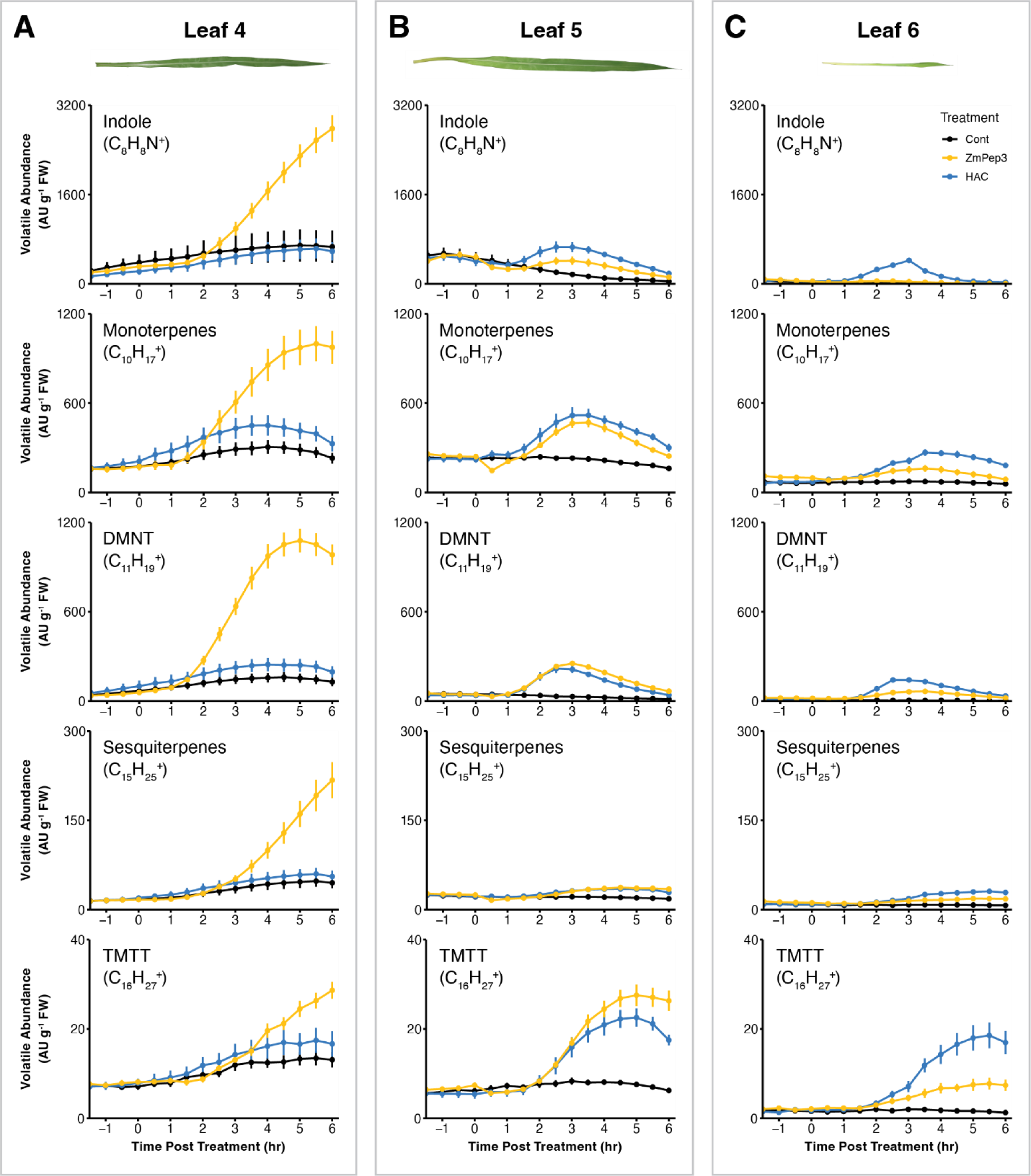
Spatial distribution of HAC-induced VOCs emission in maize. A) Volatile emissions of detached L4 leaves of V4 seedlings exposed to HAC dispensers. B) Volatile emissions of detached L5 leaves of V4 seedlings exposed to HAC dispensers. C) Volatile emissions of detached L6 leaves of V4 seedlings exposed to HAC dispensers. Leaves were either untreated (Cont), or treated with 1 µM ZmPep3, or exposed to HAC dispensers continuously. Distinguished by their exact mass, relative emission rates of indole, monoterpenes, DMNT, sesquiterpenes and TMTT were measured by PTR-MS for 6 hours after treatment. Data are shown as mean ± s. e. (n = 6).

**Supplementary Table 1.** Fold change of VOCs emission from HAC-exposed L4, L5 and L6 leaves from V4 seedlings compared to untreated leaves. Indole, monoterpenes and DMNT data were calculated based on the mean value of control leaves 3h after HAC exposure. Sesquiterpenes and TMTT data were calculated based on the mean value of control leaves 5h after HAC exposure. Data are shown as mean ± s.e. (n = 6).

| Fold Change | Indole | Monoterpenes | DMNT | Sesquiterpenes | TMTT |
| --- | --- | --- | --- | --- | --- |
| L4 | 0.80 ± 0.24 | 1.48 ± 0.24 | 1.55 ± 0.30 | 1.25 ± 0.22 | 1.25 ± 0.17 |
| L5 | 3.88 ± 0.53 | 2.24 ± 0.24 | 7.03 ± 0.76 | 1.73 ± 0.16 | 2.97 ± 0.27 |
| L6 | 19.97 ± 1.74 | 2.88 ± 0.31 | 29.53 ± 2.21 | 3.85 ± 0.30 | 11.24 ± 1.76 |

**Supplementary Figure 3.**
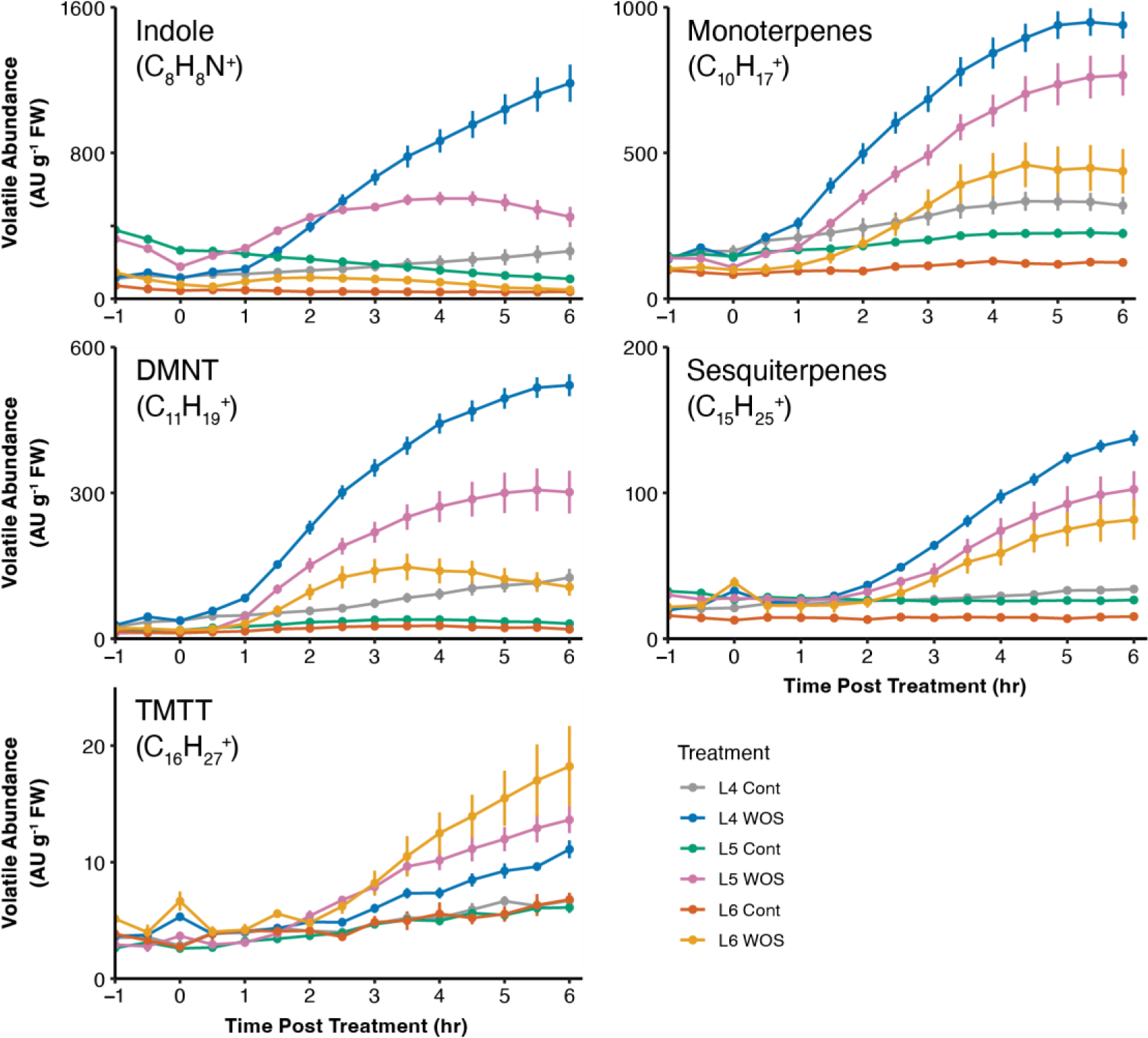
Simulated herbivory-induced VOCs emission in L4, L5 and L6 leaves of V4 seedlings. Detached L4, L5 and L6 leaves from V4 seedlings were treated by wounding and application of 10 µl *S. exigua* oral secretion (WOS). Distinguished by their exact mass, relative emission rates of indole, monoterpenes, DMNT, sesquiterpenes and TMTT were measured by PTR-MS for 6 hours after treatment. Data are shown as mean ± s. e. (n = 6).

**Supplementary Figure 4.**
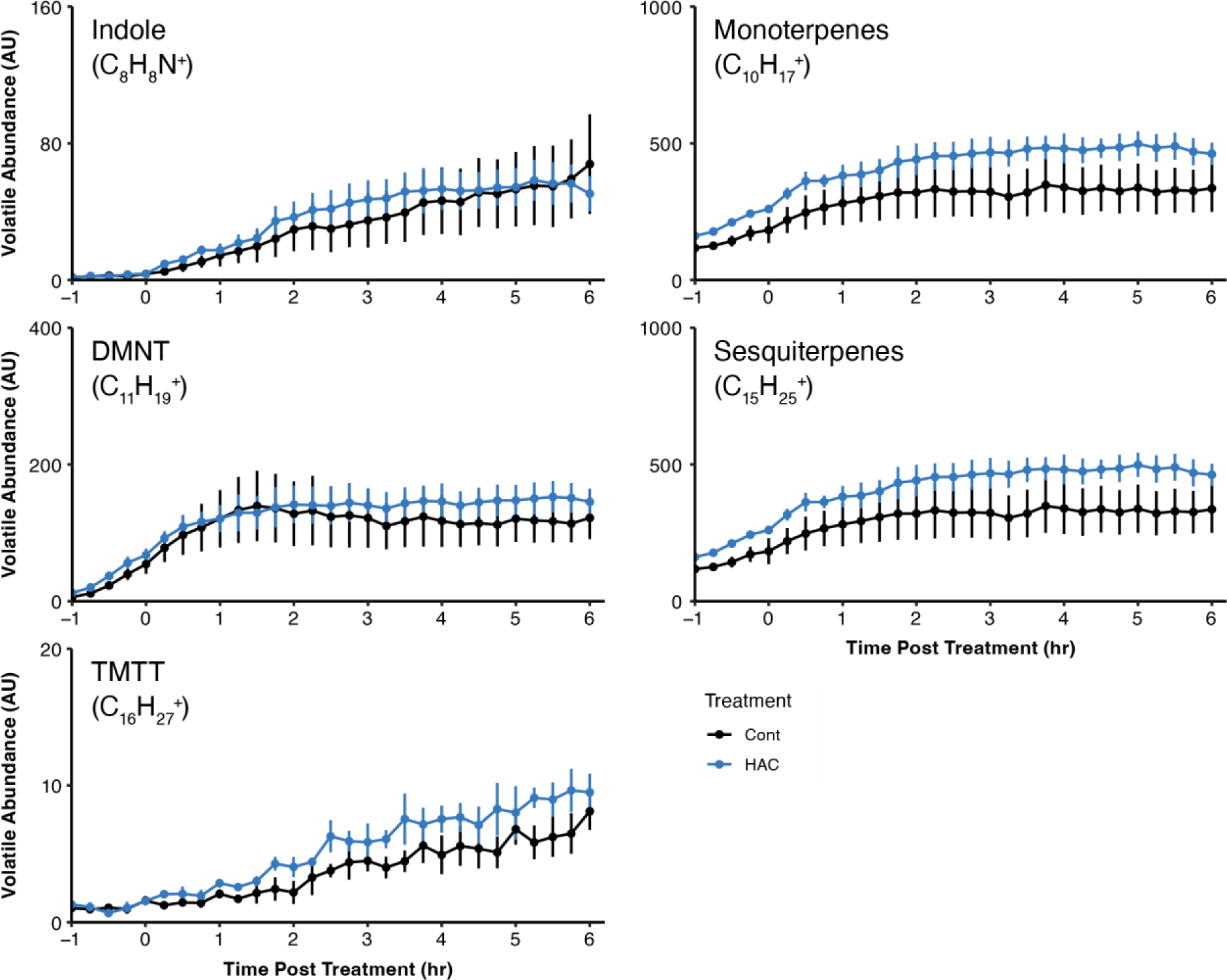
HAC-induced VOCs emission in L3 leaves of V4 seedlings. Detached L3 leaves from V4 maize seedlings were treated with HAC (HAC) dispensers. Distinguished by their exact mass, relative emission rates of indole, monoterpenes, DMNT, sesquiterpenes and TMTT were measured by PTR-MS for 6 hours after treatment. Data are shown as mean ± s. e. (n = 4).

### Volatile responsiveness to (Z)-3-hexenyl acetate is restricted to immature leaves

To determine whether the observed differences in HAC responsiveness between old and young leaves is driven by a within-leaf developmental transition, we measured HAC-induced volatile emissions in L4 leaves of different plant ages. L4 leaves of V3 seedlings (L4-V3) are expanding; L4 leaves of V4 seedling (L4-V4) are newly matured; L4 leaves of V5 seedlings (L4-V5) are mature (Supp. Fig. 5). Expanding L4 leaves (L4-V3) strongly responded to HAC exposure (Fig. 3A). Newly matured L4 leaves (L4-V4) and mature leaves L4-V5 lost their HAC responsiveness but showed enhanced responsiveness to ZmPep3 (Fig. 3B and 3C, Supp. Table 2). Thus, HAC responsiveness is a transient trait of immature maize leaves.

**Supplementary Figure 5.**
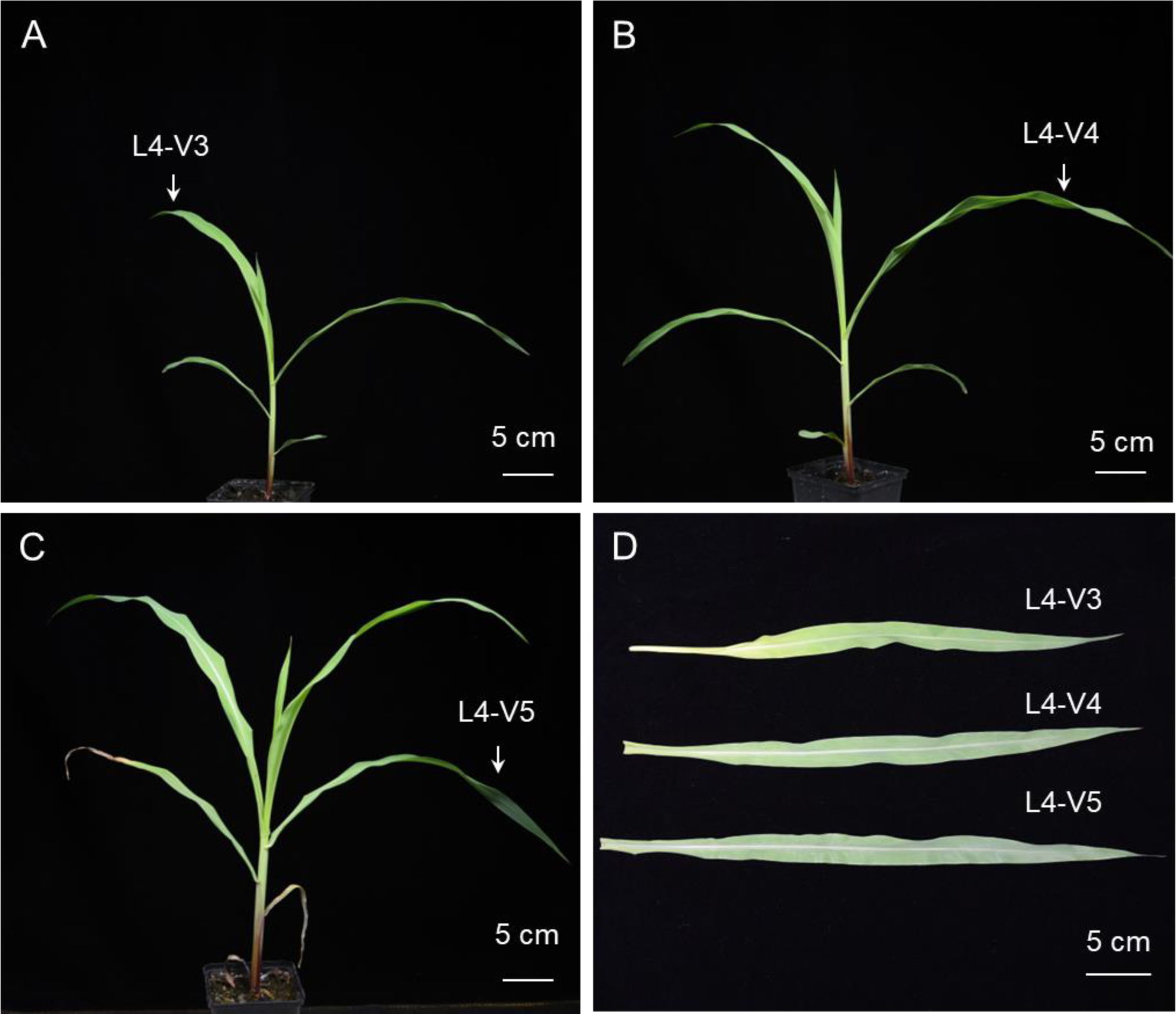
Pictures of L4 leaves from V3, V4 and V5 seedlings. A) V3 maize seedling. B) V4 seedling. C) V5 seedling. D) detached L4 from V3, V4 and V5 seedlings.

**Figure 3.**
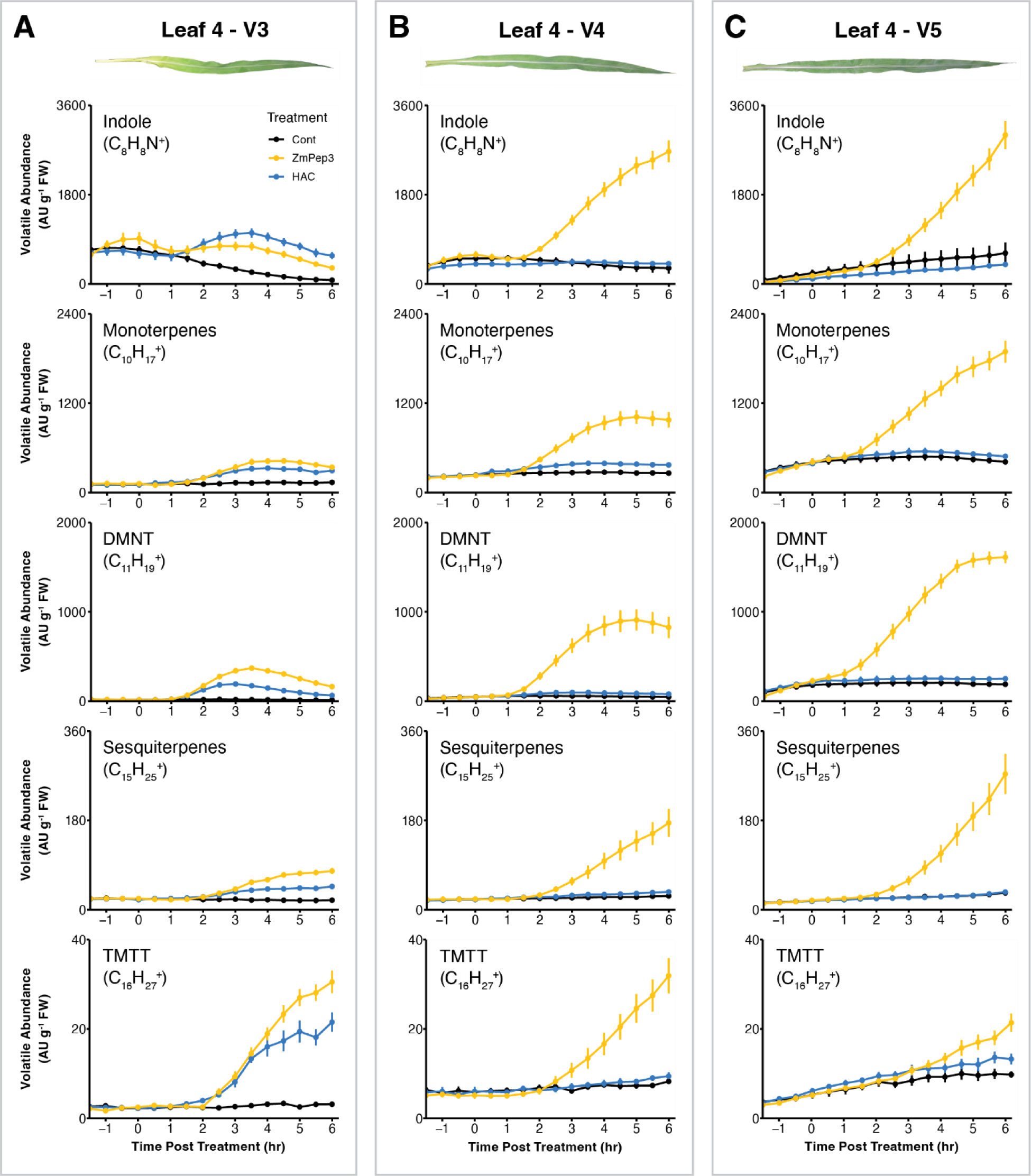
HAC-induced VOCs emission declines upon leaf maturation. A) Volatile emission of detached L4 leaves of V3 seedlings exposed to HAC (HAC) dispensers. B) Volatile emission of detached L4 leaves of V4 seedlings exposed to HAC dispensers. C) Volatile emission of detached L4 leaves of V5 seedlings exposed to HAC dispensers. Leaves were either untreated (Cont), or treated with 1 µM ZmPep3, or exposed to HAC dispensers continuously. Distinguished by their exact mass, relative emission rates of indole, monoterpenes, DMNT, sesquiterpenes and TMTT were measured by PTR-MS for 6 hours after treatment. Data are shown as mean ± s. e. (n = 6).

**Supplementary Table 2.**
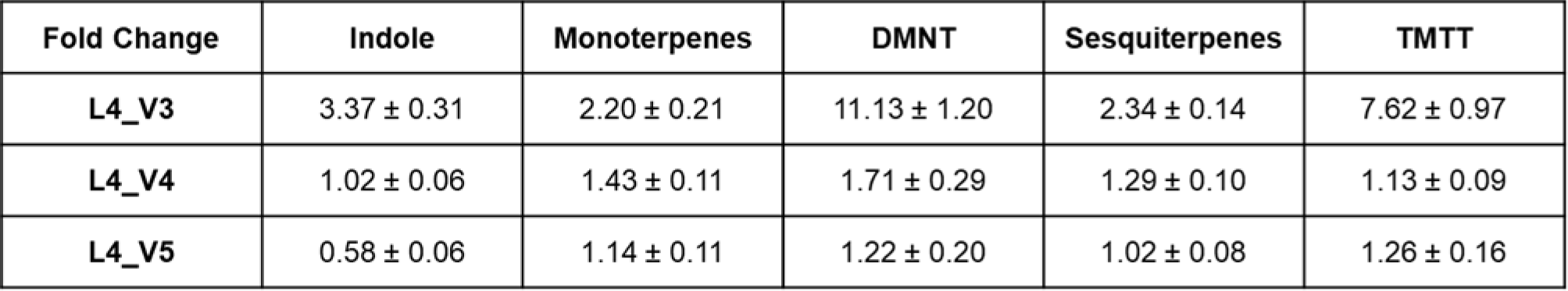
Fold change of VOCs emission from HAC-exposed L4 leaves from V3, V4 and V5 plants compared to untreated leaves. Indole, monoterpenes and DMNT data were calculated based on the mean value of control leaves 3h after HAC exposure. Sesquiterpenes and TMTT data were calculated based on the mean value of control leaves 5h after HAC exposure. Data are shown as mean ± s.e. (n = 6).

### Volatile responsiveness to the full blend of herbivory induced plant volatiles is restricted to immature leaves

Plants are typically exposed to complex blends of herbivory induced plant volatiles (HIPVs). To test if volatile responsiveness to HAC is also reflected in responsiveness to entire volatile blends, we exposed L4 and L6 leaves from V4 seedlings to volatiles of non-attacked maize plants and maize plants that were infested with *Spodopera exigua* caterpillars. We exposed the leaves to one of the two volatile blends overnight, then supplied these leaves with clean airflow for 2 hours, after which point we measured volatile emission. Compared to exposure of L6 to volatiles of control plants, exposure of L6 to volatiles of *S. exigua* infested plants resulted in enhanced emissions of monoterpenes, sesquiterpenes and TMTT. Green leaf volatiles, indole and DMNT emissions were unchanged. By contrast, we found no significant difference in volatile emission for L4 leaves (Fig.4). Thus, volatile responsiveness to full blends of herbivory induced plant volatiles is restricted to younger maize leaves.

**Figure 4.**
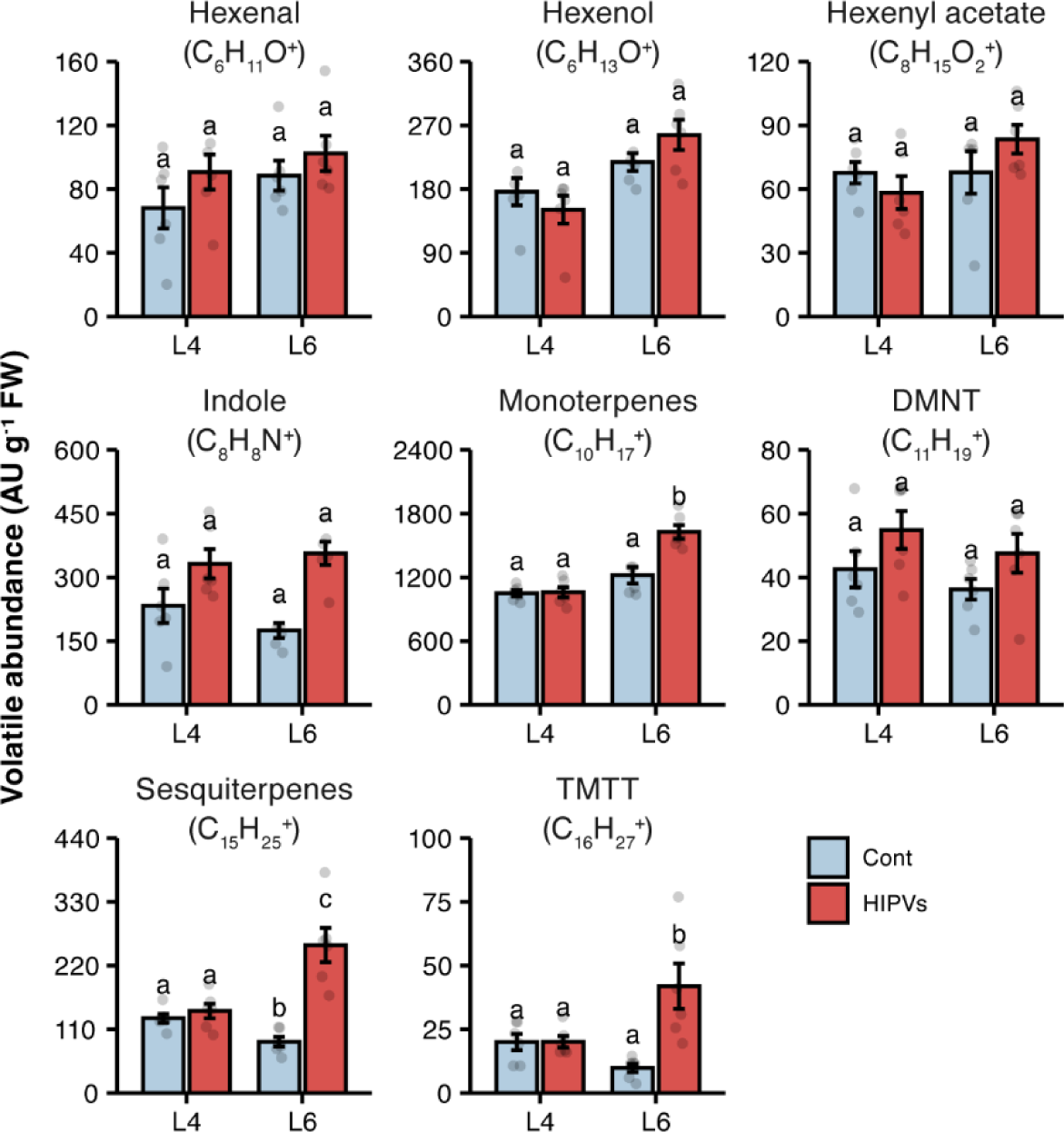
Immature leaves are the major sensing organs for HIPVs. L4 or L6 leaves from V4 seedlings were exposed to the headspace volatile from uninfested (Cont) or *S.exigua* infested maize plants (HIPVs) for 10 hours and were further incubated in clean air for 2 hours (1 hour in dark, then 1 hour under light), after which point relative emission rates of hexenal, hexenol, hexenyl acetate, indole, monoterpenes, DMNT, sesquiterpenes and TMTT were determined by PTR-MS. Different letters above the bars indicate statistical significance at P < 0.05. Data are shown as mean ± s. e. (n = 6).

### Leaf cuticular structure does not determine volatile responsiveness to (*Z*)-3-hexenyl acetate

One major change during maize leaf maturation is the altered wax type and chain length of the cuticle(Bourgault et al., 2020). Together with the accumulation of cutin, these changes lead to the formation of a water-proof cuticle. As cuticular composition may alter volatile uptake and release, it may drive ontogenetic changes in volatile responsiveness (Camacho-Coronel et al., 2020; Liao et al., 2021; Wang and Erb, 2022). To test if the composition of the leaf cuticle affects HAC responsiveness, we compared HAC-induced volatile emission of L4, L5, and L6 leaves of V4 seedlings between wild type and the *glossy6* mutant plants. The *glossy6* mutant has reduced epicuticular wax load compared to the wild type (Li et al., 2019). By consequence, its leaves are “leaky” and less water resistant (Supp. Fig. 6). Overall, we observed lower constitutive and induced volatile levels in the glossy6 mutant than B73 wild type plants (Fig. 5A). However, the HAC and ZmPep3 responsiveness pattern was largely unchanged in the mutant: L4 leaves were unresponsive to HAC, but highly responsive to ZmPep3 treatment (Fig. 5A, Supp. Fig. 7). L5 and L6 leaves on the other hand responded strongly to HAC, but less to ZmPep3 (Fig. 5B and 5C, Supp. Fig. 7). An exception was TMTT, which was more strongly induced by ZmPep3 in L5 leaves than L4 leaves. These results show that the leaf cuticular wax composition, and, by inference, leaf cuticular development, is not the driving factor that determines HAC responsiveness across leaf ontogeny.

**Supplementary Figure 6.**
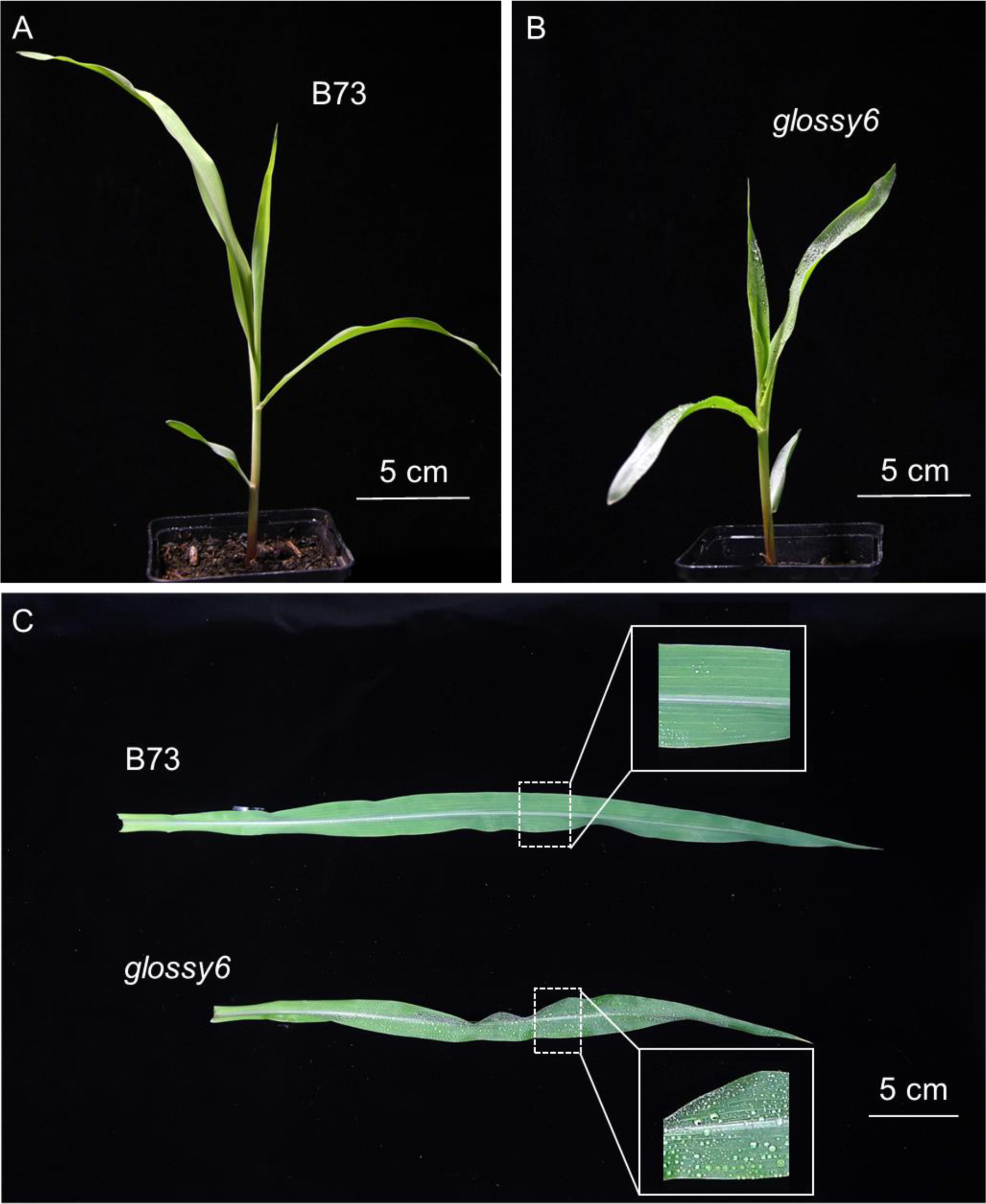
Pictures of *glossy6* mutants. A) V2 stage B73 seedling (wild type). B) V2 stage *glossy6* seedling. C) Detached L4 leaves from V4 stage B73 and *glossy6* plants. Enlarged area shows the adaxial side of leaf blades after water spraying.

**Figure 5.**
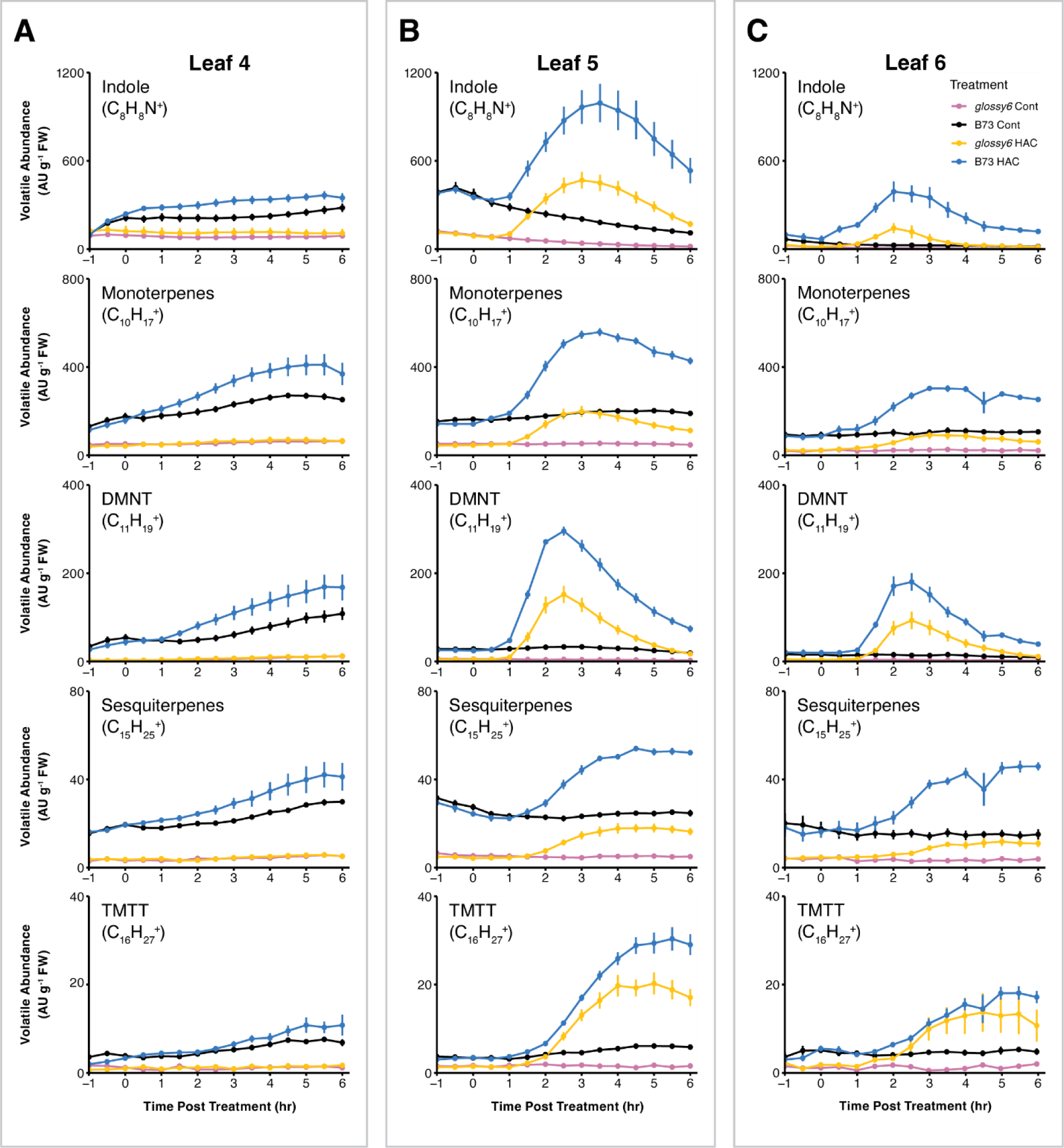
Leaf cuticular structure does not determine ontogenetic patterns of HAC responsiveness. A) Volatile emission of detached L4 leaves from B73 or *glossy6* V4 seedling exposed to HAC dispensers. B) Volatile emission of detached L5 leaves from B73 or *glossy6* V4 seedlings exposed to HAC dispensers. C) Volatile emission of detached L6 leaves from B73 or *glossy6* V4 seedlings exposed to HAC dispensers. Leaves were either untreated (Cont), or exposed to HAC dispensers continuously. Distinguished by their exact mass, relative emission rates of indole, monoterpenes, DMNT, sesquiterpenes and TMTT were measured by PTR-MS for 6 hours after treatment. Data are shown as mean ± s. e. (n = 6).

**Supplementary Figure 7.**
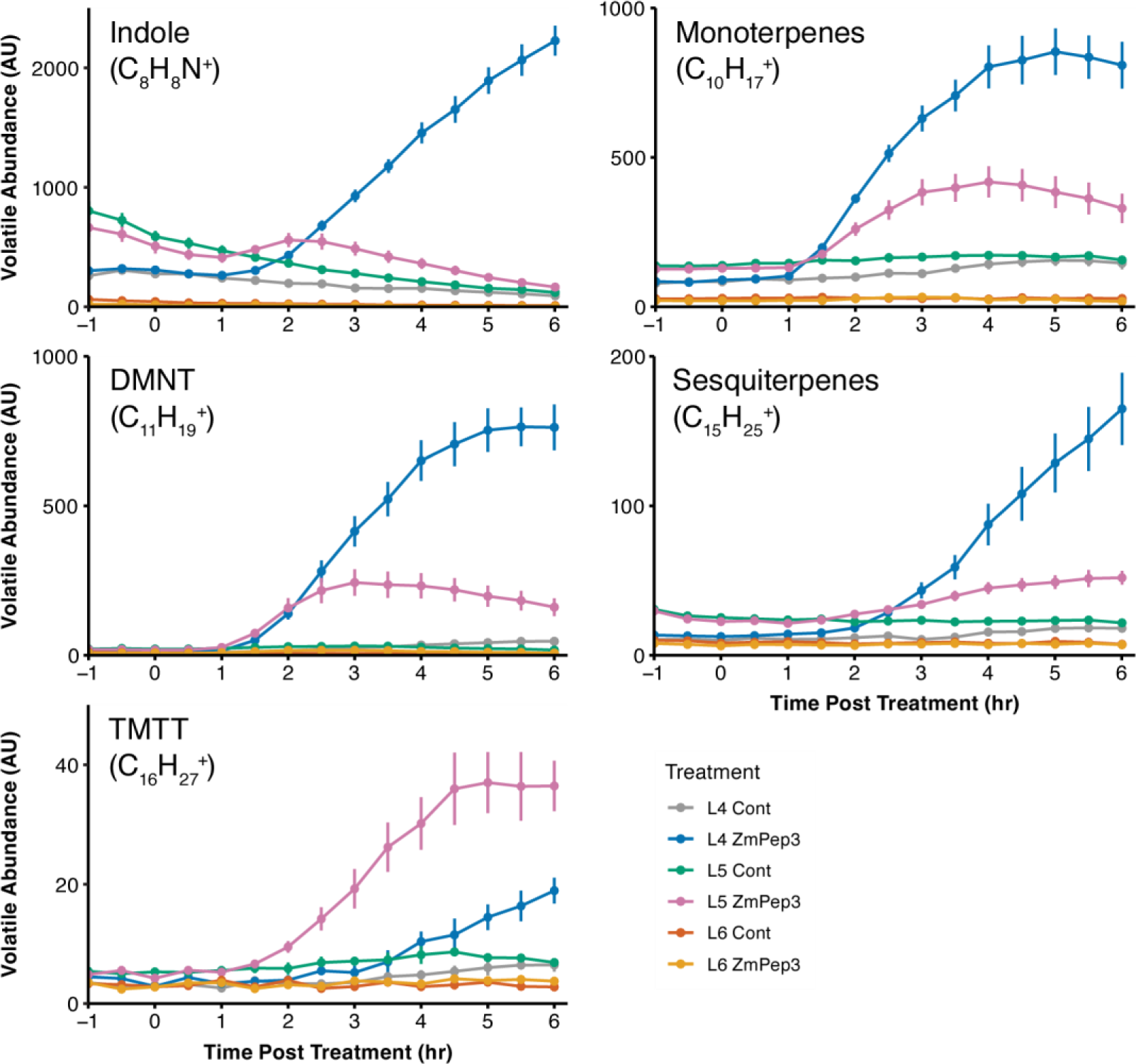
ZmPep3-induced VOCs emission in leaves of the *glossy6* plants. Detached L4, L5 and L6 leaves from V4 stage *glossy6* mutants were either untreated (Cont) or treated with 1 µM ZmPep3. Distinguished by their exact mass, relative emission rates of indole, monoterpenes, DMNT, sesquiterpenes and TMTT were measured by PTR-MS for 6 hours after treatment. Data are shown as mean ± s. e. (n = 6).

### Gas exchange via stomata does not match responsiveness to (*Z*)-3-hexenyl acetate

Stomata are seen as the main exchange point of volatiles between the environment and the leaf apoplast. Thus, we evaluated gas exchange rate of leaves at various developmental stages to test whether HAC responsiveness of immature leaves may be explained by higher stomatal conductance (Wang et al., 2019; Kong et al., 2021). Within a maize seedling, stomatal conductance is lower in the youngest L6 leaves than the older L4 and L5 leaves (Supp. Fig. 8A). Within a given leaf, gas exchange increases as the leaf matures and stomata are fully developed (Supp. Fig. 8B). If stomatal function and gas exchange were the driving factor explaining HAC responsiveness, older and mature leaves should respond more strongly to HAC. As we observe the opposite pattern, gas exchange via stomata is not the major driver that explains the transient responsiveness of immature maize leaves to HAC.

**Supplementary Figure 8.**
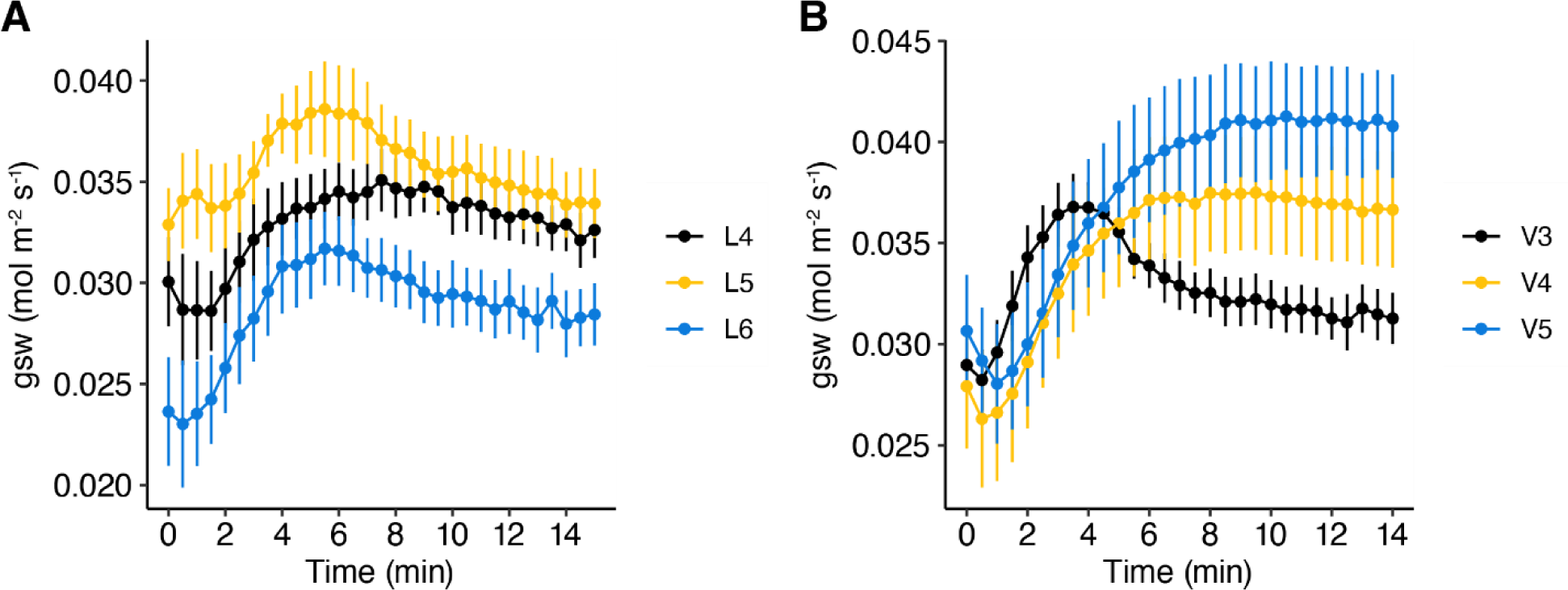
Stomatal conductance of various maize leaves. A) Absolute stomatal conductance (g_sw_) of L4, L5 and L6 leaves from V4 stage B73 seedlings. B) Absolute stomatal conductance (g_sw_) of L4 leaves from V3, V4 and V5 stage B73 seedlings. Data are shown as mean ± s. e. (n = 5).

### Transcriptional reprogramming by (Z)-3-hexenyl acetate is strongest in young leaves

To test whether the strong, specific HAC responsiveness of young leaves is a general pattern or whether it is constrained to volatile induction and release, we analyzed transcriptomic responses of L4, L5 and L6 leaves to HAC exposure. We selected two time points, 0.5 h and 2 h after HAC exposure, to evaluate the transcriptional change before and after the onset of HAC-induced volatile emission. We also measured ZmPep3 responses 2 h after ZmPep3 treatment for comparison. 0.5 h after HAC exposure, there were 3313 and 2405 differentially expressed genes (DEGs) in L5 and L6 leaves, respectively. L4 leaves only had 1492 DEGs (Fig. 6A, Supp. Fig. 9). The difference in DEGs was even more pronounced 2 h after HAC exposure. There were 4458 DEGs in L5 and 5860 DEGs in L6, and only 568 DEGs in L4 leaves (Fig. 6A). In contrast, ZmPep3 modulated the expression of a similar number of genes across all tested leaves (Fig. 6A). We also observed less overlap between HAC regulated genes than ZmPep3 regulated genes when comparing different leaves (Fig. 6A), demonstrating that there is a strong and specific interaction between HAC responsiveness and leaf age. At the same time, this experiment shows that L4 leaves still retain residual HAC responsiveness.

To further understand the HAC-induced regulation of defensive processes in young and old leaves, we focused our transcriptional analysis on JA biosynthesis genes, JA signaling genes, volatile and benzoxazinoid biosynthesis genes (Hu et al., 2018; Block et al., 2019; Erb and Reymond, 2019; Wang et al., 2020). In accordance with the volatile emission patterns (Fig. 2, Supp. Table 1), induction of the majority of the volatile biosynthesis genes by HAC was significantly higher in L5 and L6 leaves than L4 leaves (Fig. 6B). Two terpene synthases genes, *TPS5* and *TPS23*, showed a higher fold change in L4 leaves. Closer inspection revealed that this pattern was due to a very low basal expression level in L4 leaves rather than higher expression upon HAC treatment (Supp. Fig. 10). ZmPep3-triggered induction of volatile biosynthesis genes was similar across the different leaf stages.

The expression of JA biosynthesis and signaling genes followed the same pattern as the volatile biosynthesis genes, with HAC inducing a stronger response in L5 and L6 leaves than L4 leaves, and ZmPep3 triggering similar up-regulation among the three leaf stages (Fig. 6C). The expression of *LOX8*, *AOS1a*, *AOS1c*, *JAR1a*, *MYC7*, *JAZ5*, *JAZ6*, *JAZ8*, and *CYP94B3a* was rapidly up regulated upon HAC treatment (Fig. 6C and 6D). LOX8, AOS1a, AOS1c are enzymes critical for the biosynthesis of 12-OPDA. JAR1 is responsible for the conjugation of isoleucine to JA to form the active JA-Ile. MYC7 is the maize ortholog of Arabidopsis MYC2—the master regulator of the JA signaling cascade. JAZ proteins are JA transcription repressor and coreceptor. CYP94B3a is critical for JA catabolism (Borrego and Kolomiets, 2016; Howe et al., 2018). Their rapid induction illustrates that JA signaling is an early event in HAC triggered defense (Engelberth et al., 2004). The finding that they are more strongly induced in L5 and L6 than L4 leaves illustrates that HAC responsiveness in mature leaves is impaired upstream of jasmonate biosynthesis and signaling.

Benzoxazinoids have a central role in maize chemical defense and are regulated by jasmonate signaling (Zhou et al., 2018). Several benzoxazinoid biosynthesis genes were strongly induced by HAC. Again, HAC induction was much higher in L5 and L6 than L4 leaves. Responsiveness to ZmPep3 was comparable across leaves (Fig. 6E). Thus, differential sensitivity to HAC also translates into changes in the transcriptional expression of direct defense metabolite pathways.

**Figure 6.**
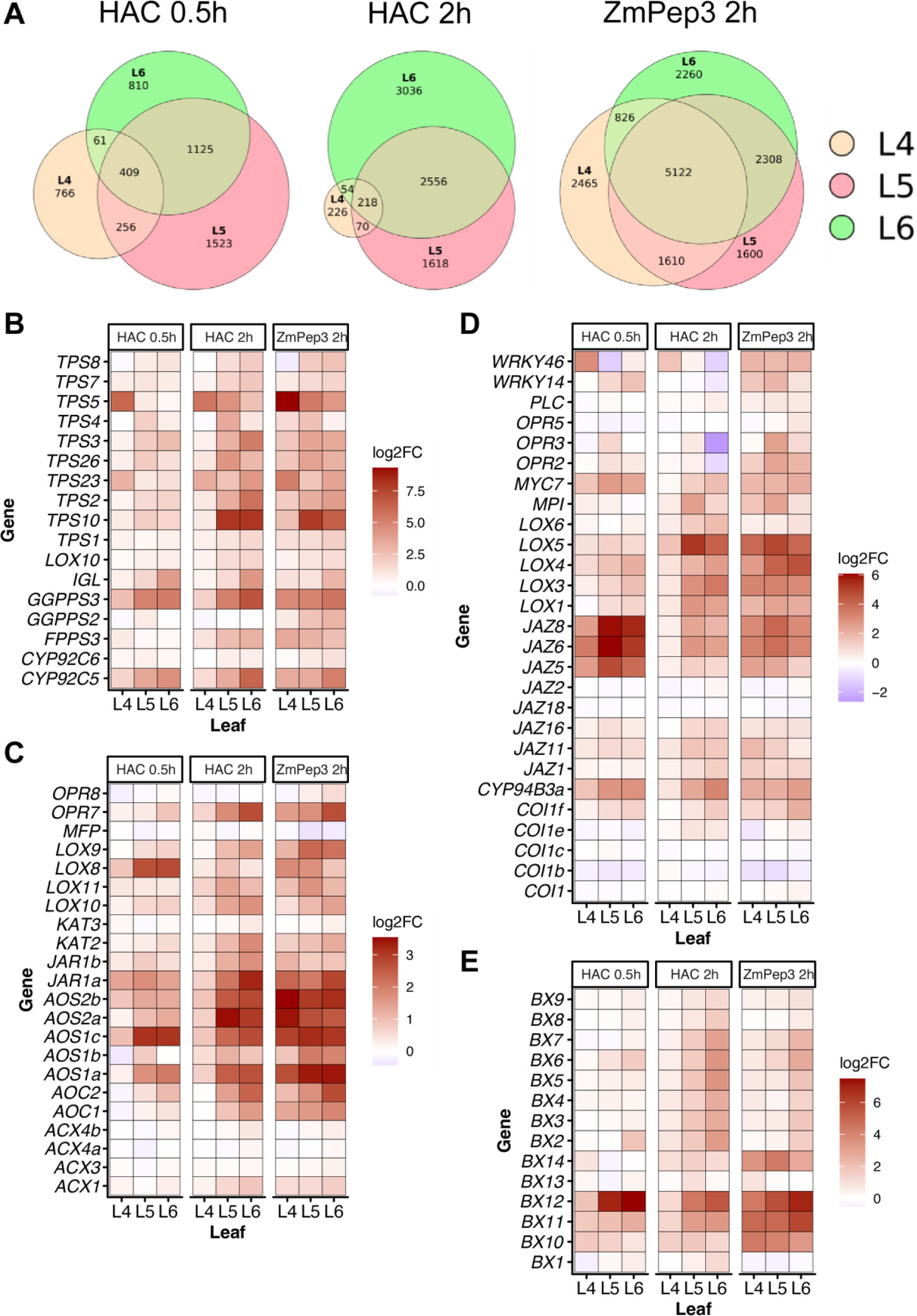
HAC-induced transcriptional reprogramming declines with leaf maturation. A) Venn diagram of the DEGs in L4, L5 and L6 leaves after HAC 0.5h, HAC 2h or 1 µM ZmPep3 2h treatment. B) Fold change of the expression of volatile biosynthesis genes. C) Fold change of the expression of JA biosynthesis genes. D) Fold change of the expression of JA signaling genes. E) Fold change of the expression of benzoxazinoids biosynthesis genes. Fold change of gene expression were analyzed based on log2(FPKM Value+1)-transformed data. All leaves X treatments samples were prepared with 6 biological replicates.

**Supplementary Figure 9.**
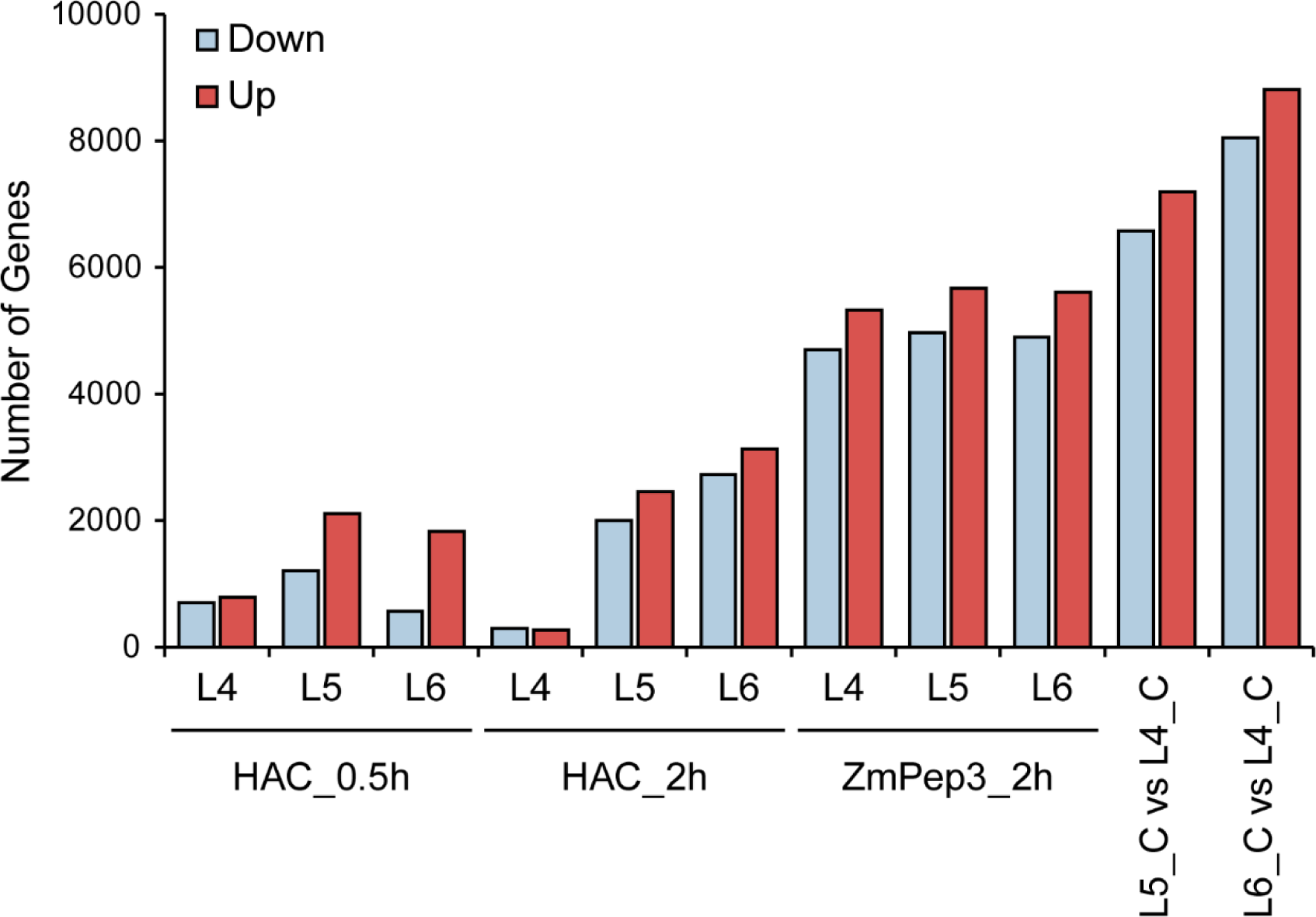
Number of differentially expressed genes in HAC or ZmPep3 treated maize leaves.

**Supplementary Figure 10.**
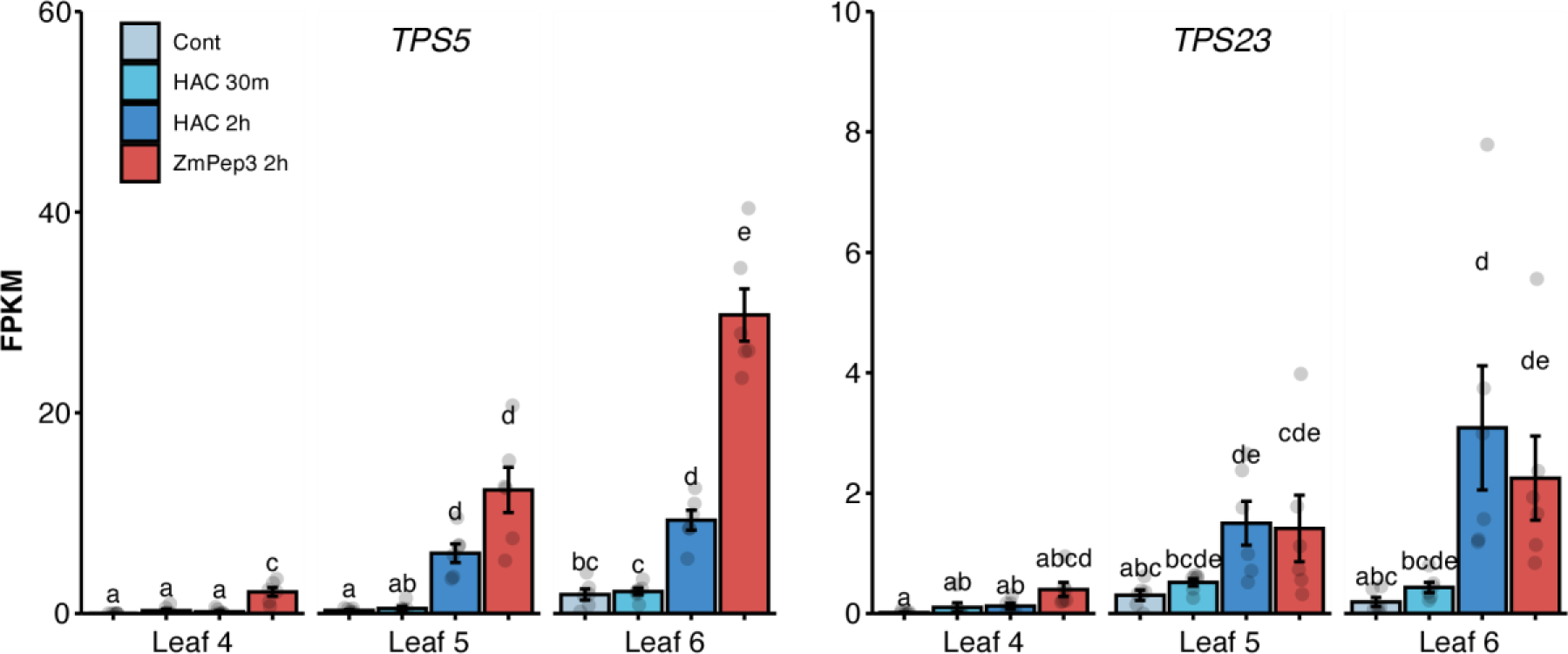
Reads of the *TPS5* and *TPS23* transcripts in HAC or ZmPep3 treated maize leaves. Different letters above the bars indicate statistical significance at P < 0.05. Data are shown as mean ± s. e. (n = 6).

### Jasmonate induction by (Z)-3-hexenyl acetate is strongest in young leaves

Jasmonates regulate volatile and non-volatile secondary metabolite biosynthesis in maize. To test whether the different induction of jasmonate biosynthesis genes by HAC also translates into different jasmonate levels, we measured jasmonate accumulation in HAC and ZmPep3 treated L4, L5 and L6 leaves. HAC rapidly induced the accumulation of JA and JA-Ile (Fig. 7). Both basal and HAC-induced levels of JA and JA-Ile were markedly higher in L5 and L6 than L4 leaves 0.5 h after HAC exposure. After 2 hours of HAC exposure, JA and JA-Ile concentration in L4 leaves dropped back to basal levels but remained elevated in L5 and L6 leaves. OPDA levels remained unchanged after HAC exposure (Fig. 7). ZmPep3 increased JA and JA-Ile contents across all leaves. Induction of JA was highest in L6, followed by L5 and L4. JA-Ile was highest in L4, followed by L6 and L5. Together with the transcriptomic data, these data show that HAC induces high sustained levels of jasmonates in young, but not in old leaves.

**Figure 7.**
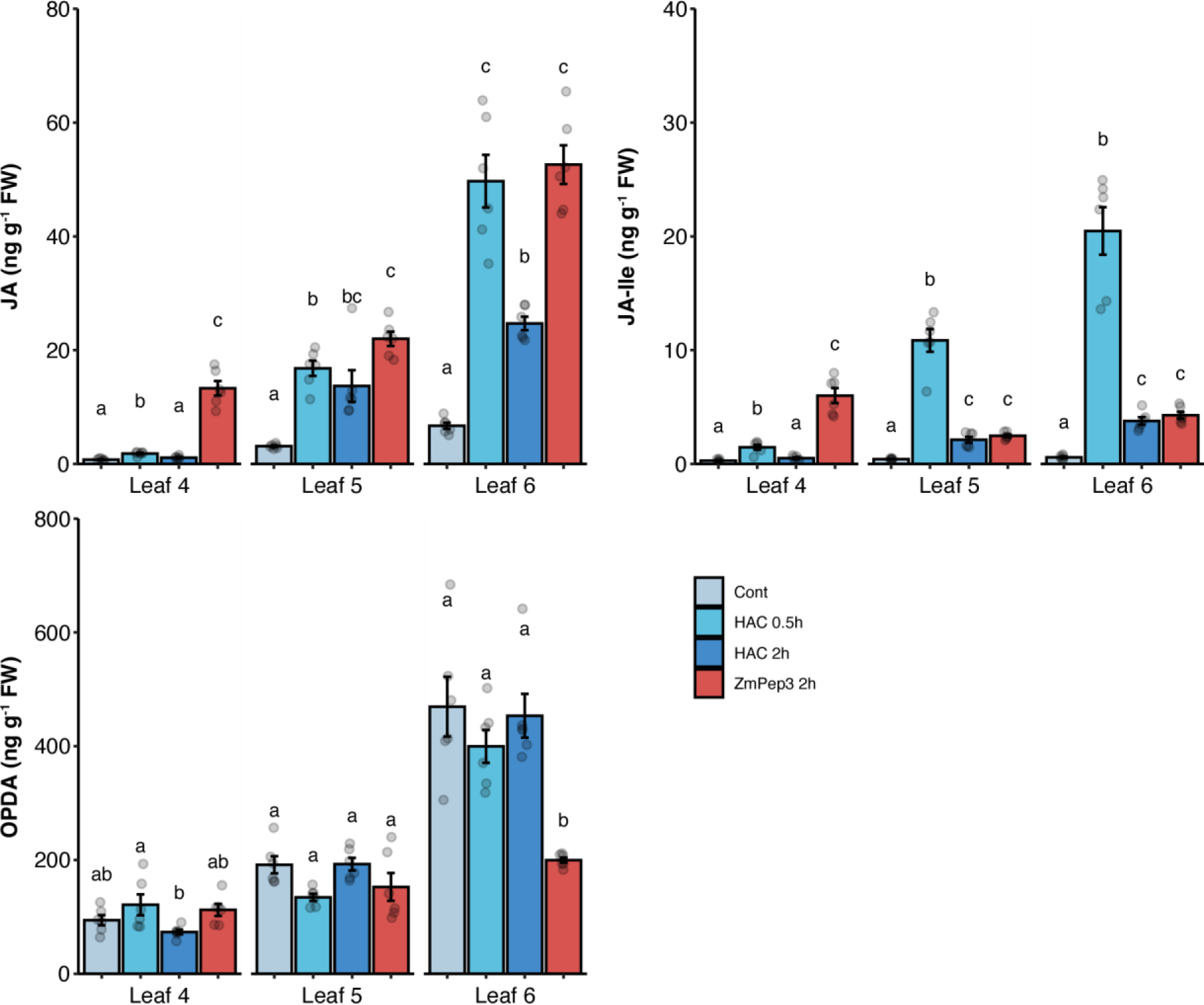
HAC induces lower and less sustained JA and JA-Ile accumulation in mature leaves. JA, JA-Ile and OPDA contents in L4, L5 and L6 leaves after HAC 0.5h, HAC 2h or 1 µM ZmPep3 2h treatment. Statistical significance was calculated within each leave types. Different numbers above the bars indicate statistical significance at P < 0.05. Data are shown as mean ± s. e. (n = 6).

## Discussion

Plants can detect herbivory-induced volatiles as danger cues. The perception of stress volatiles triggers the activation of defenses, which helps plants to fend off incoming herbivores. So far, it remains largely unknown which shoot tissues are capable of sensing plant volatiles, and how this sensitivity changes as plants grow and develop (Wang and Erb, 2022). We uncover that maize plants perceive herbivory-induced volatiles such as HAC predominantly with their young, immature leaves. As soon as leaves mature, they lose their capacity to respond strongly to HAC, even though their canonical defense signaling machinery is fully functional and highly sensitive to non-volatile defense cues such as the peptide ZmPep3. We thus propose that young leaves serve as transient volatile sensing organs in maize.

Leaf development includes several distinct stages with different functional profiles. Changes between stages can be rapid, a phenomenon referred to as heteroblasty (Zotz et al., 2011). In grasses for instance, young developing leaves are photosynthetically much less active than old leaves, as their stomata are not fully formed yet (Lawson and Blatt, 2014; Nunes et al., 2020; Kong et al., 2021). Conversely, leaf movement in Mimosa plants is strong in young leaves, but cedes in old leaves (Allen, 1969). Here, we show that young maize leaves are highly sensitive to to herbivory induced volatiles such as HAC. This responsiveness drops rapidly as leaves mature, and fully developed leaves are hyposmic, showing only very weak responsiveness to HAC at the transcriptional, hormonal and volatile level. This leads to a spatial patterning of volatile perception where young leaves act as transient volatile sensing organs and lose this capacity as they mature. The capacity of maize plants to perceive volatiles thus dynamically decreases from plant tip to base. It is tempting to speculate that plants with more complex architectures may have hundreds of volatile-sensitive leaves that are constantly replaced by new leaves as the plants grow. Akin to root tips exploring the rhizosphere (O’Brien et al., 2016), this could result in a perceptive mosaic where the outermost regions of the shoots are most capable of exploring the volatile landscape that the plants grow into.

What is the mechanism behind the stronger responsiveness of young maize leaves to volatiles? One possibility is that defense signaling is increasingly impaired in old leaves, thus making those leaves generally less responsive to defense elicitors (Zhang et al., 2020). Our experiments with ZmPep3 show that this is not the case. On the contrary, mature leaves respond more strongly to this non-volatile early defense elicitor, from jasmonate signaling to downstream defense and volatile deployment. Another possibility is that physiological differences such as different cuticular composition or stomatal behavior may impair volatile uptake of younger leaves (Sugimoto et al., 2016; Liao et al., 2021; Wang and Erb, 2022). Our experiment with the cuticular *glossy6* mutant indicates that cuticular composition is not a major determining factor of volatile responsiveness. However, we cannot fully rule out that physical differences in leaf surface or cellular permeability may contribute to the observed phenomenon. Stomatal density and gas exchange are lower rather than higher in young leaves (Kong et al., 2021), making impaired uptake through stomata in mature leaves unlikely. Given these considerations and the fact that HAC unresponsiveness is evident already at the early signaling level, we hypothesize that the molecular mechanisms that are required for HAC perception or early signaling transduction are preferentially expressed in young maize leaves and are largely inactivated in older leaves. Comparable patterns are known for other sensing mechanisms. In *Nicotiana benthamiana* for instance, the immune receptor CORE for sensing CSPs shows an age-dependent expression pattern, with low expression in young plants and higher expression in older plants. This expression pattern leads to the CSPs-insensitive young plants gaining CSPs responsiveness as they age (Wang et al., 2016). In the case of volatile sensing, the rapid shift in HAC sensitivity will prove useful to identify early signaling events, including potential receptors involved in HAC perception.

Is there an adaptive or agricultural benefit to constraining volatile perception to young leaves? More experiments will be needed to answer this question, but our results together with the current literature provide a number of potential benefits of this behavior. Many plants grow and compete within dense canopies. Thus, most of the mature leaves are embedded in crowded canopies, where short-range diffusion of volatiles is enhanced, but long-range transfer may be less effective. Thus, placing volatile sensing organs on top of the canopy may benefit long range sensing. Another advantage may be found in herbivore feeding patterns and within-plant volatile signaling. In maize for instance, generalist lepidopteran herbivores start feeding on old leaves and then move to younger leaves (Köhler et al., 2015). Thus, having mature leaves that are sensitive to herbivory and emit large amounts of volatiles together with emerging leaves that are highly sensitive to volatiles may favor beneficial within-plant signaling, where plants can effectively protect their young emerging leaves via the rapid systemic induction of defenses.

In conclusion, this work shows that young maize leaves are highly responsive to stress volatiles, and rapidly lose this responsiveness as they mature. The transient nature of a plant’s “nose” has important consequences for the spatiotemporal perception patterns of plant volatiles, the capacity of plants to sense volatiles at different positions within a canopy and the preferred routes of systemic defense signaling within plants. Our work provides a basis for a deeper mechanistic understanding of plant volatile perception and the spatiotemporal mosaic of plant volatile interactions in plant canopies.

## Material and Methods

### Plant material

Maize (*Zea mays*, line B73) plants were used throughout this study. The *glossy6* mutants (B73 background) were kindly provided by Patrick Schnable (Iowa State University). Maize seeds were sown directly into pots (9 cm x 9 cm x 10 cm) with commercial soil (Selmaterra, BiglerSamen, Switzerland). All plants were grown in a glasshouse supplemented with artificial lights (∼300 μmol m^-2^ s^-1^) at 22 ± 2 °C, 40–60% relative humidity (RH), with a 14 h: 10 h, light: dark cycle.

### HAC dispensers

HAC dispensers were prepared as previously described with minor modifications (Hu et al., 2019). In brief, 1.5-ml glass vials (Ø×H 11.6 × 32 mm) containing ∼ 100 mg glass wool were filled with 200 µl (*Z*)-3-hexenyl acetate (HAC, >98%, Sigma-Aldrich, Buchs, Switzerland) and sealed with screw caps containing a rubber septum. The caps were then pierced with a 1-µl glass capillary (Drummond) and sealed with PTFE tape. Finished vials were wrapped with aluminum foil and equilibrated for at least 2 days before using. Dispensers with approximately 70 ng*h^-1^ HAC emission, which corresponds to the GLVs emission rate from herbivore attacked maize, were selected for experiments.

### Synthetic peptides

ZmPep3 (>90%) was synthesized by GenScript and dissolved in sterile Milli-Q water as 10 mM stock before use.

### Plant treatments

V4 seedlings were cut at the position of 2^nd^ leaf with a razor blade. For experiments with detached seedlings, L3 leaves were carefully removed, and the remaining seedlings were further cut by 0.5 cm under Milli-Q water to remove air-filled xylem to facilitate water uptake. For experiments with detached leaves, L4, L5 and L6 leaves were carefully separated, all leaves were further cut by 0.5 cm under water. For L4 leaves (mature leaves), an additional removal of leaf sheath was required before further cutting of the leaf blade under water. Leaves from V3 and V5 seedlings were prepared similarly, except that the first cut was done at the L1 position for V3 seedlings and at the L3 position for V5 seedlings. All plant materials were transferred into glass beakers (250 mL) with 150 mL Milli-Q water and allowed for recovery for at least 2 hours before treatment.

Detached leaves or seedlings were transferred into 12-ml plastic centrifuge tubes containing 10 ml Milli-Q water. These samples were then transferred into the glass cylinders (see “Volatile Measurement” for details) for at least 2 hours acclimation, during which the background volatile emission was monitored. ZmPep3 treatment was done by adding 10 μl of a 1mM ZmPep3 solution in the centrifuge tubes to achieve a final concentration of around 1 µM. HAC treatment was done using the volatile dispensers (as described above). Simulated herbivory was achieved by wounding the leaf blades 3 times (0.5 cm with a hemostatic forcep) on each side of the mid-vein on the leaf blades and supplying 10 µl *S. exigua* oral secretions with a pipette. HIPVs exposure was done by connecting the airflow outlet of a glass cylinder containing a detached V4 seedling infested by 5 4-instar *S.exigua* larvae to the airflow inlet of a glass cylinder containing an experimental leaf. For volatile measurements, ZmPep3 and HAC treatments were started immediately after the background volatile measurements. Simulated herbivory was carried out 5 minutes before the start of the volatile profiling. HIPVs exposure was done for 10 hours (2 hours under light, 8 hours in dark) before disconnecting the emitters and receivers, and reconnecting the receivers with clean air flow for 2 h (1 hour in dark, 1 hour under light), after which point volatile emissions were determined.

### Volatile measurements

Plant volatile profiling was conducted with an automated high-throughput volatile sampling platform. This platform consists of a proton transfer reaction time-of-flight mass spectrometer (PTR-TOF-MS) system (Tofwerk, Switzerland), an automated headspace sampling system (Abon Life Sciences, Switzerland), and 100 transparent glass cylinders (Ø×H 12 × 45 cm, with transparent lids) supplied with clean airflow (0.8 L min^-1^) through an airflow inlet and outlet placed on benchtop above the PTR-TOF-MS. The PTR-TOF-MS system draws air at 0.1 L min^-1^, ionizes the contained VOCs using protonated H_2_O and analyzes them in real time.

Volatiles emitted by plants inside the glass cylinders were purged by clean airflow to the airflow outlets, where the PTR-TOF-MS sampling inlet can be cycled using the fully automated programming routines of the sampling system. Each round of volatile sampling took 15 minutes or 30 minutes, and every sample took 14 seconds or 27 seconds, respectively. Between samples, a fast zero air measurement was performed for 3 seconds to minimize contamination. Complete mass spectra (0-500 m/z) were recorded in positive mode at ∼10 Hz and averaged to a single point per sample. The PTR was operated at a temperature of 100 C and an E/N of approximately 120 Td. The volatile data extraction and processing were then done with the Tofware software package v3.2.2 (Tofwerk, Switzerland).

Illumination for the plants was provided by LED lights (DYNA, heliospectra) placed ∼ 80 cm above the glass cylinders to provide light at ∼300 μmol m^-2^ s^-1^ during volatile sampling.

### Leaf-level gas exchange measurements

Gas exchange measurements in the glasshouse were done with a LI-6800 Portable Photosynthesis System (Li-COR Biosciences Inc., Lincoln, NE, USA) equipped with a Multiphase Flash Fluorometer (6800-01A) chamber. The middle position of the expanded leaf blade was measured for 15 minutes using the 2 cm^2^ leaf chamber. Conditions in the LI-6800 chamber were as follows: flow rate, 500 μmol s^-1^; fan speed, 10000 rpm; leaf temperature, 30 °C; RH, 40%; [CO2], 400 μmol mol^-1^; photosynthetic active radiation (PAR), 1800 μmol m^-2^ s^-1^.

### Transcriptome analysis

Leaf samples for transcriptome analysis were prepared as described in plant treatment and volatile measurement. Whole leaf samples were harvested with 6 biological replicates for each leaf type/treatment/time combination by flash freezing in liquid nitrogen. Total RNA extraction was done with the GeneJET PCR Purification Kit (K0702, Thermo Scientific). DNAse I (M0303, New England BioLabs) treatment was done to minimize genomic DNA contamination. Total RNA sample quality control was done with the Agilent 2100 Bioanalyzer (Agilent RNA 6000 Nano Kit). Library construction was done by BGI (Shenzhen, China) and the transcriptome sequencing was done with the BGISEQ platform.

Bioinformatics were done by BGI. In brief, reads of low quality, adaptor sequence or high levels of N base were filtered to get clean reads. The resulting clean reads were aligned to the maize genome (GCF_000005005.2_B73_RefGen_v4) with HISAT.

### Phytohormone profiling

Leaf samples for phytohormone profiling were prepared the same way as for transcriptome analysis. Phytohormone profiling were done with an ultra-high-pressure liquid chromatography-tandem mass spectrometry (UHPLC/MS-MS) as previously described (Glauser et al., 2014). In brief, around 100 mg of fine leaf powder were extracted with 990 μL EtOAc: formic acid, 99.5:0.5 (v/v) and 10 μL internal standard solution containing isotopically labelled hormones (d_5_-JA, ^13^C_6_- JA-Ile, d_6_-ABA, d_6_-SA, d_2_-GA3, and d_5_-IAA, 1μg/mL each in 70% MeOH).The mixture was centrifuged at 14,000 × g for 3 min and the supernatant was collected into a 2 mL centrifuge tube. The pellet was extracted again with 500 μL EtOAc: formic acid, 99.5:0.5 (v/v). After repeating the previous procedure, the supernatants were combined and evaporated till dryness. The residual was resuspended in 100 μL 70% MeOH. After centrifugation, 5 μL were injected into the UHPLC/MS-MS. The hormones were quantified by calculating a calibration equation obtained by linear regression from five calibration points for each compound. Peak areas of the hormones measured in the samples were normalized to the internal standard before applying the calibration equation. Finally, hormone levels were normalized to the weight of plant sample used.

### Statistical analyses

Differences in *TPS5* expression were analyzed using a two-way ANOVA (type II) on log+1-transformed data. A value of 1 was added to each value as some expression values, pre-transformation, were 0. *TPS23* data did not meet the assumptions of ANOVA, and therefore the non-parametric Scheirer-Ray-Hare test was used. Differences in phytohormone concentrations between treatments in L4, L5 and L6 leaves were, where possible, analyzed using one-way ANOVAs, and data were log-transformed when untransformed data were not normally distributed. For JA in leaf 6 and OPDA in leaf 4, data were analyzed using heteroscedasticity-consistent standard errors with White-adjusted ANOVAs (White, 1980). Differences in JA for leaf 5 were analyzed using a Kruskal-Wallis test as the data did not meet the assumptions of ANOVA following transformation. The effect of HIPVs from *S. exigua*-infested plants on plant VOCs emission and whether this was leaf age-dependent was tested using two-way ANOVA at the individual volatile level. When the interaction between HIPV exposure and leaf age was significant, we conducted Tukey’s HSD tests for pairwise comparisons. All statistical analyses were conducted in R version 4.1.1. All ANOVAs were run using the R package ‘rcompanion’ (Mangiofico, 2022).

## Author contributions

L.W. and M.E. conceived and supervised the project. L.W. and M.E. designed experiments. L.W., S.J., T.C. and M.W. performed experiments. L.W., T.C. and J.W. analyzed data and created figures. L.W. and M.E. wrote the manuscript with input from all authors.

## Acknowledgements

We thank the laboratory of Patrick Schnable (Iowa State University) for providing *glossy6* mutant seeds. We thank Gaetan Glauser (University of Neuchâtel) for assistance with phytohormone measurement. We also thank members of the whole Biotic Interactions group for helpful discussions.

